# Bone Morphogenetic Protein Pathway Modulates Parkinson’s Disease Genetic Risk and Promotes Motor Recovery

**DOI:** 10.64898/2026.06.08.730818

**Authors:** Aleksandar Rajkovic, Zhao Zhang, Rakesh Sahu, Lang Liu, Mahima Mishra, Dimitrii Komkov, Ilan Rosenstein, Joy Kahn, Roland H. Friedel, Ziv Gan-Or, Claude Brodski

**Affiliations:** Department of Physiology and Cell Biology, Zelman Center for Neuroscience, Faculty of Health Sciences, Ben-Gurion University of the Negev, 84105 Beer Sheva, Israel; Montreal Neurological Institute, McGill University, Montreal, Quebec, Canada H3A 1A1; Department of Human Genetics, McGill University, Montreal, Quebec, Canada H3A 1Y2; Department of Neurology and Neurosurgery, McGill University, Montreal, Quebec, Canada H3A 2B4; Nash Family Department of Neuroscience, Friedman Brain Institute, Icahn School of Medicine at Mount Sinai, New York, USA

**Keywords:** Parkinson’s disease, polygenic risk score (PRS), Genome wide association study (GWAS), α-synuclein, bone morphogenetic proteins (BMP), gene therapy, neurotrophic factors

## Abstract

Progress toward disease-modifying Parkinson’s disease (PD) therapies is hindered by limited understanding of pathogenic drivers and lack of therapies that restore the damaged dopaminergic (DA) neurons driving motor deficits. We investigated the role of the bone morphogenetic protein (BMP) pathway in PD using common and rare genetic variants across large-scale datasets. Single-variant analyses identified nominal associations with PD risk and onset age. A BMP polygenic risk score (PRS) was significantly associated with PD risk (OR = 1.21, empirical P = 1.0 × 10⁻⁴), a result replicated in proxy-case analyses (OR = 1.14, empirical P = 1.0 × 10⁻⁴) and remained significant after sensitivity testing. To test the result’s functional relevance, we genetically and pharmacologically inhibited BMP signaling in mice, which induced motor deficits and PD-like neuropathology. In a α-synuclein preformed fibril (PFF) mouse model, BMP5 and BMP7 (BMP5/7) demonstrated neuroprotective effects when administered *concurrently* with α-synuclein PFFs. Importantly, when delivered *after* PFFs-induced motor symptoms onset, BMP5/7 demonstrated neurorestorative effects, ameliorating both motor impairments and neuropathology. These findings provide first evidence that BMP signaling variations contribute to polygenic PD risk, identify a physiological role for this pathway in safeguarding against PD-related pathology, and demonstrate BMP’s therapeutic potential for disease modification.

## Introduction

Parkinson’s disease (PD) is a neurodegenerative disorder characterized by progressive motor impairments affecting over 10 million people worldwide, with currently no disease-modifying therapy available that can stop or slow the disease course ^1, 2^. The motor symptoms are caused by the loss of dopaminergic (DA) neurons of the substantia nigra pars compacta (SNpc). α-synuclein-associated neuropathological changes are thought to play a central role in the degeneration of DA neurons ^3, 4^, with phosphorylated α-synuclein (p129), serving as a standard marker for PD neuropathology ^5^.

By the time PD is diagnosed by motor symptom onset, α-synuclein pathology and DA degeneration are already widespread in the brain ^6–8^. Therefore, the time between diagnosis and the extensive depletion of remaining DA neurons causing debilitating motor impairments has been regarded as a critical time window of opportunity for disease-modifying therapies ^9^.

The etiology of PD is complex, including genetic, epigenetic, and environmental factors ^10, 11^. Familial PD accounts for up to 15% of the total cases ^12^. Most of the disease is considered to be sporadic, likely caused by the cumulative effect of multiple common or rare variants with small to moderate effect sizes, in addition to other unknown environmental and stochastic factors. Large-scale genome-wide association studies (GWAS) have substantially advanced our understanding of the genetic architecture of idiopathic PD, identifying up to 134 independent risk loci to date ^13^. Despite these important discoveries, most associated variants individually confer only modest effects on disease risk, and the causal genes and underlying biological mechanisms at many loci remain unclear ^14^. Functional interpretation of genetic variants provides a critical opportunity to bridge this gap by linking risk loci to specific biological pathways and cellular processes, helping to elucidate molecular mechanisms and identify potential therapeutic targets for PD.

Neurotrophic factors have been suggested as prime candidates for disease modification in PD due to their strong survival-promoting effects on neurons ^15–17^. Some of these proteins have been tested in clinical studies for PD ^18^, including the most widely tested substance, glial cell-line-derived neurotrophic factor (GDNF), as well as neurturin (NRTN), and cerebral dopamine neurotrophic factor (CNTF). While all were shown to be generally safe, they did not meet their primary endpoint ^16^. It has been suspected that insufficient distribution of neurotrophic factors played an important role in the disappointing outcomes of clinical trials. Therefore, significant improvements in delivery techniques in recent years have encouraged further clinical trials testing GDNF, with very promising initial results ^19^.

However, there are concerns that neurotrophic factors are not efficient in rescuing neurons affected by α-synuclein-associated pathology when applied at the time of diagnosis. While there are reports demonstrating the neuroprotective effects of neurotrophic factors when administered *together* with α-synuclein, the neurorestorative effects of these proteins when given *after* α-synuclein have not been established ^16, 20^. Therefore, studying the neurorestorative effects of neurotrophic factors in α-synuclein-based PD models is considered a critical step in further validating neurotrophic factors as therapeutic candidates for PD ^16^.

Bone morphogenetic proteins (BMPs) are neurotrophic factors that play a central role in a wide array of cellular processes. The canonical BMP pathway is initiated by the binding of BMP ligands to their receptors. This triggers the intracellular activation of SMAD1/5/8 effectors, which then complex with SMAD4 to regulate gene transcription, while inhibitory SMAD6/7 provide essential negative feedback. This signaling cascade is further modulated by extracellular antagonists and intracellular SMURFs, which also participate in other independent signaling networks ^21^.

BMP5 together with BMP7 (BMP5/7) are essential regulators of the development of DA neurons *in vivo,* signaling through SMAD1 to control the differentiation of DA progenitors, particularly to SNpc neurons ^22^. Moreover, BMP5/7 increase the maturation of human stem cells to DA neurons ^22^. BMPs are not only essential for the development of DA neurons but also have neuroprotective effects in toxin-based PD models ^23^. Moreover, BMP5/7 and BMP14 (also known as growth differentiation factor 5) showed neuroprotective effects in viral α-synuclein overexpression models when applied simultaneously with α-synuclein ^24, 25^. Mice in which the SMAD1 gene was inactivated in embryonic stem cells that can differentiate to neurons or glia showed an accumulation of α-synuclein in adulthood, suggesting that the BMP pathway can affect α-synuclein homeostasis ^25^. However, if BMPs can exert neurorestorative effects in already damaged DA neurons, and if inactivation of BMP signaling would lead to PD-specific motor and neural pathology, remain unknown.

In our current study, we investigated whether BMP pathway abnormalities can contribute to PD risk and age at onset (AAO) by performing human genetics studies. To assess the functional relevance of these findings, we inactivated BMP signaling genetically specifically in differentiated DA neurons and pharmacologically in adult mice and studied PD-associated motor and neuropathological changes. Finally, we used the α-synuclein PFFs model to assess the neuroprotective effects of BMP5/7 in comparison to GDNF and the neurorestorative properties when applied to damaged neurons after the onset of motor symptoms.

## Results

### Common and rare variant analyses implicate BMP pathway polygenic risk in PD

We first evaluated the association between common variants (MAF > 0.01) within the BMP pathway and PD risk, as well as age at onset (AAO), using genotyping data from the Canadian Open Parkinson Network (C-OPN) and the UK Biobank (UKB; 3,597 cases and 35,970 controls) and regression-based models. To capture potentially undiagnosed PD cases and individuals at earlier disease stages, we repeated the PD risk analysis with proxy cases included (18,611 cases and 186,110 controls). Several variants showed nominal associations with PD risk and AAO. However, none remained significant after correction for multiple testing (Supplementary Table S1 – S3).

To assess the cumulative contribution of common variants, we further performed pathway-specific polygenic risk score (PRS) analyses. The BMP pathway PRS showed a significant association with PD risk (OR = 1.21, empirical P = 1.00 × 10⁻⁴), but not with AAO (OR = 0.01, empirical P = 9.94 × 10⁻^1^; Table 1). This association was replicated in analyses including proxy cases (OR = 1.14, empirical P = 1.0 × 10⁻⁴). To assess the robustness of these findings, we further evaluated the pathway after removing genes encoding extracellular BMP antagonists, SMURFs, or both. Although these regulators are essential to BMP signaling, they are also significantly involved in several other pathways ^26–31^. All analyses remained significant (Supplementary Tables S4–S5).

**Table 1.** Association of BMP pathway PRS with PD risk and AAO. BMP pathway PRS is strongly associated with PD risk in analyses with and without proxy cases and shows a modest association with AAO only in proxy-expanded analyses.

| Outcome | Proxy Cases | Effect Size | Num_SNP | P-value | Empirical P |
| --- | --- | --- | --- | --- | --- |
| PD risk | No | OR = 1.21 | 1508 | $3.73 \times 10^{-25}$ | $1.00 \times 10^{-4}$ |
| | Yes | OR = 1.14 | 1499 | $2.24 \times 10^{-64}$ | $1.00 \times 10^{-4}$ |
| AAO | No | $\beta$ = 0.01 | 1202 | $9.25 \times 10^{-1}$ | $9.94 \times 10^{-1}$ |

OR, odds ratio; $\beta$ , beta; P, p-value; **Empirical P**, permutation-derived p-value; **Num\_SNP**, number of SNPs.

To evaluate the contribution of rare variants (MAF < 0.01), we conducted pathway-based burden analyses using SKAT-O and meta-analysis approaches using whole genome sequencing data from AMP-PD (3,051 cases and 3,667 controls) and UKB (without proxy: 3,173 cases and 31,722 controls; with proxy: 18,110 cases and 180,962 controls). In the UKB proxy cohort, a nominally significant association was detected for deleterious (CADD-filtered) variants (P = 0.03). However, this signal was not retained in sensitivity analyses (Supplementary Tables S6–S7).

### Conditional deletion of SMAD1 in postmitotic embryonic DA neurons induces PD-associated motor impairments and neuropathology in adult mice

To validate the functional and biological relevance of our findings, we investigated how disruptions to the canonical BMP signaling pathway affect DA neurons. To do so, we deleted SMAD1 in these neurons and studied PD-associated pathologies. We bred *Smad1^fl/fl^* mice in which the start codon of the *Smad1* gene is flanked by loxP sites ^32, 33^ with *Dat^Cre^* knock-in mice which have Cre recombinase expression directed to DA neurons, without disrupting endogenous dopamine transporter expression (Fig. 1A) ^34^. This approach allowed us to directly assess the contribution of SMAD1 to the maintenance of motor function and α-synuclein homeostasis in differentiated DA neurons, as DAT expression starts with DA neuronal differentiation.

**Figure 1.**
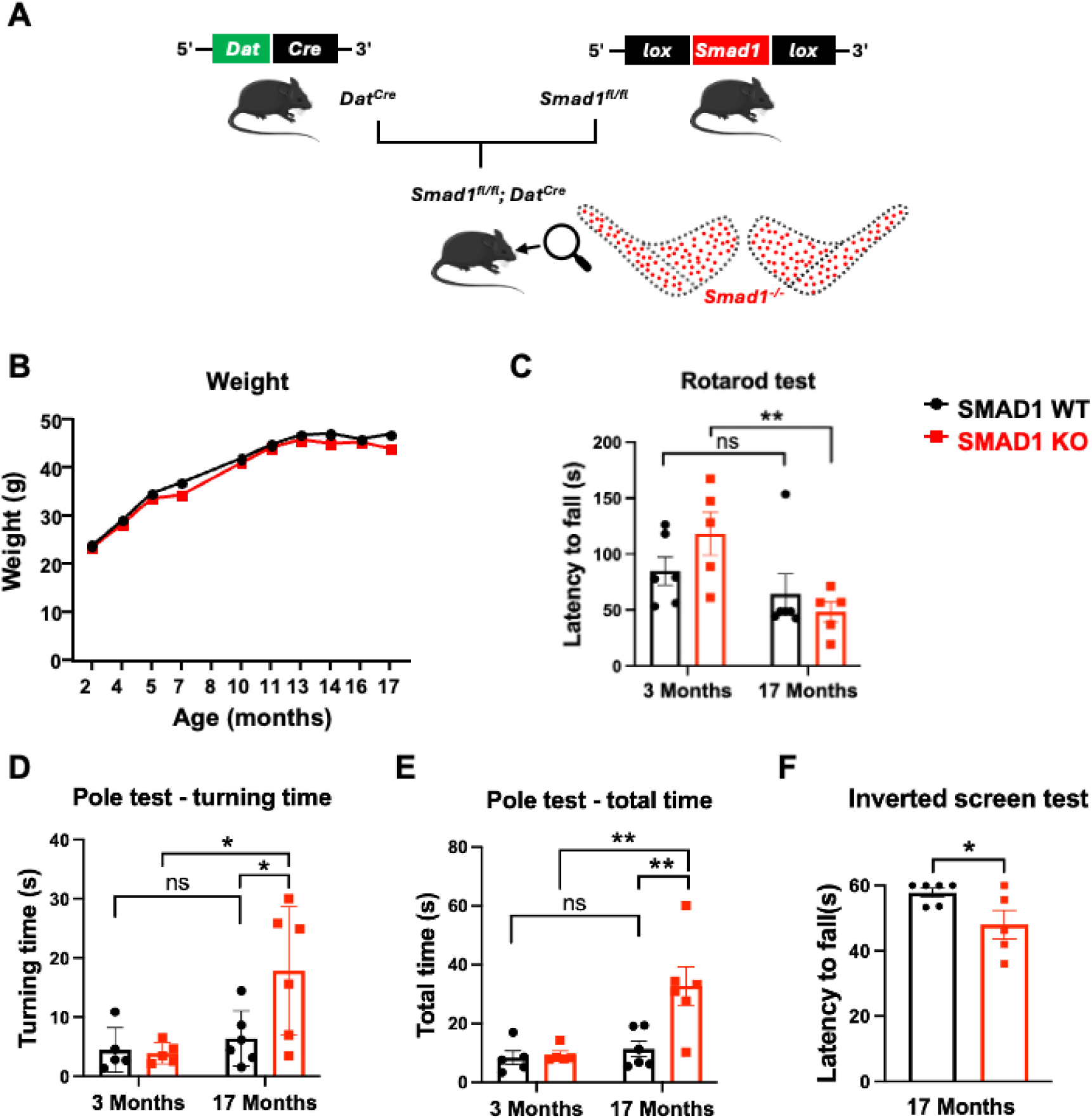
Conditional SMAD1 KO mouse mutants show progressive motor deficits. **(A)** Breeding scheme for the generation of *Smad1^fl/fl^;Dat^Cre^* (SMAD1 KO), lacking SMAD1 in postmitotic DA neurons. **(B)** Body weight of SMAD1 conditional knockout (KO) and control animals (WT) over 17 months indicates no significant differences between genotypes. **(C)** Rotarod performance in young (3-month-old) and late middle-aged (17-month-old) animals, revealing a significant longitudinal decline in latency to fall exclusively within the SMAD1 KO cohort. **(D)** Pole test turning time, defined as the latency to orient vertically downward, demonstrates intact early-stage agility but a significant, age-dependent performance decay in 17-month-old SMAD1 KO mutants. **(E)** Pole test total time required to completely descend the pole indicated an increased descent duration occurring in 17-month-old SMAD1 KO mice. **(F)** Muscle strength and endurance were reduced in 17-month-old mutants as measured by the latency to fall in the inverted screen test. Data in (C–E) were analyzed using a two-way ANOVA with a mixed-effects model, followed by Šídák’s post hoc multiple-comparison test to evaluate aging trajectories within each genotype. Data in (F) were analyzed using an unpaired t-test. Significance levels are indicated as follows: *p < 0.05; **p < 0.01; ns = non-significant.

First, we used conditional SMAD1 knockout (*Smad1^fl/fl^; Dat^Cre^*) and littermate wild-type (WT) control mice to evaluate the effects of SMAD1 deficiency on age-related motor changes. To do this, we assessed the motor behavior of young adult (3-month-old) and late middle-aged (17-month-old) mice. Throughout the time we followed the animals, we did not observe changes in weight between controls and mutants (Fig. 1B). In the rotarod test, mutants exhibiting a significant age-dependent decline in latency to fall between 3 months and 17 months of age (predicted means 118.5 s vs. 48.27 s; adjusted p = 0.0050), while WT mice showed no significant change over the same interval (predicted means 85.06 s vs. 64.31 s; adjusted p = 0.3806; main effect of age: F(1, 9) = 15.69, p = 0.0033; Fig. 1C). In the pole test, significant differences in turning time between WT and SMAD1-deficient mice were present at 17 months of age (predicted means 6.39 s vs. 17.80 s; adjusted p = 0.0145), with no genotype difference detected at young baseline (adjusted p = 0.9890), indicating worsening motor coordination and agility in the mutant group (F(1, 8) = 8.104, p = 0.0216; Fig. 1D). Consistently, total pole descent time revealed significant genotype differences at the aged endpoint (predicted means 11.37 s in WT vs. 32.75 s in SMAD1-deficient mice; adjusted p = 0.0021), but not at young baseline (adjusted p = 0.9812), supported by a significant age-by-genotype interaction (F(1, 8) = 6.283, p = 0.0366; Fig. 1E). Because the inverted screen test was implemented later in the project, we did not evaluate 3-month-old wild-type (WT) and mutant animals. However, at 17 months of age, SMAD1-deficient mice exhibited a significantly decreased latency to fall compared to controls (Fig. 1F)

The neuropathological analysis of 17-month-old animals revealed significant abnormalities in SMAD1-deficient mice. In the striatum, mutants showed reduced striatal TH fluorescence intensity, indicating a loss of dopaminergic projections (Fig. 2A, B). Additionally, there was a significant increase in mutants in both the total area and mean fluorescence intensity of pS129-a-syn in this brain area, indicating pathological α-synuclein accumulation (Fig. 2A, C, D). In the SNpc, no significant differences were observed between genotypes regarding the TH-positive area or the count of dopaminergic neurons (identified by NeuN+/TH+ co-staining) (Fig. 2E, F, G). However, in mutants total p129-α-synuclein area was significantly increased in the SNpc as well as the percentage of TH+ area containing p129-α-synuclein expression (Fig. 2E, H, I).

**Figure 2.**
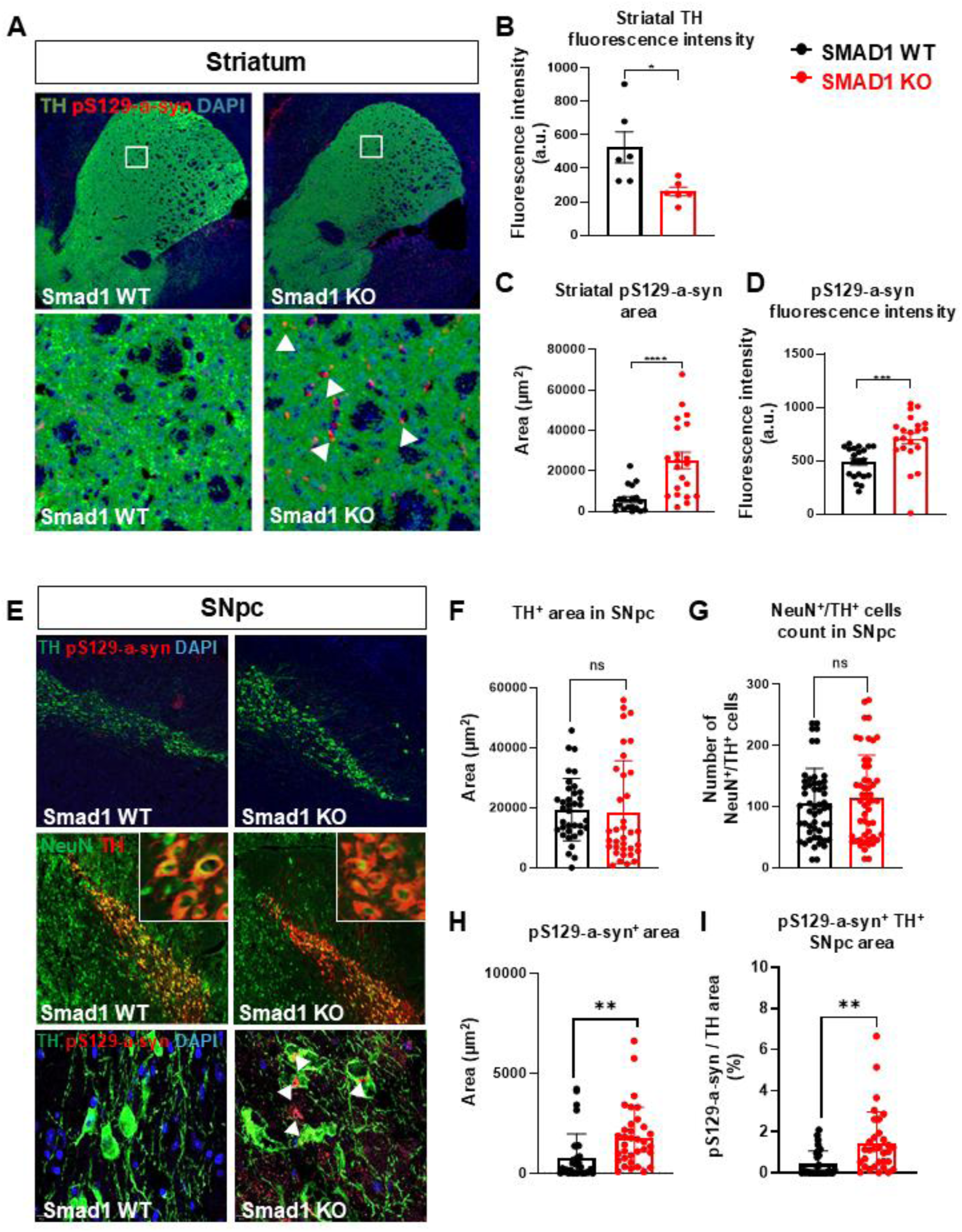
Conditional SMAD1 KO mouse mutants exhibit PD-associated neuropathology. **(A)** Immunostaining of striatal TH and p129-α-synuclein in WT and SMAD1 KO. **(B)** Quantification of striatal TH fluorescence intensity, indicate reduced dopaminergic terminal integrity in SMAD1 KO. **(C, D)** p129-α-synuclein area and intensity is increase in SMAD1 KO mutants. **(E)** Immunostaining of nigral TH and p129-α-synuclein. **(F, I)** SNpc TH area and DA neuronal numbers (identified by NeuN+/TH+ co-staining) are not different between the genotypes. **(G, H)** p129-α-synucelin area are increased in the SNpc and in the TH+ area. Statistical analysis was conducted using unpaired t-tests. Significance levels are indicated as follows: *p < 0.05; **p < 0.01; ***p < 0.001.

Together, we conclude that SMAD1 plays an essential role in differentiated DA neurons for the maintenance of adult motor function and α-synuclein homeostasis.

### Pharmacological blockade of BMP/SMAD signaling in adult mice by LDN-212854 causes PD-associated motor impairments and neuropathological deficits

We then aimed to determine the consequences of BMP/SMAD signaling inhibition in adult mice. To do this, we pharmacologically inactivated BMP/SMAD signaling in 6-month-old mice using the selective small-molecule BMP receptor inhibitor LDN-212854 ^35^. After a 7-day training period in a behavioral test battery, animals were injected intraperitoneally with either LDN-212854 or vehicle for 14 days as previously shown to effectively block BMP signaling (Fig. 3A) ^35^. During the treatment period, treated animals showed a significant reduction in their weight (Fig. 3B). In the rotarod test, LDN-212854-exposed mice showed a significantly reduced latency to fall (Fig. 3C). Moreover, in the pole test, treated mice exhibited prolonged turning time and total descending time (Fig. 3D, E). In the inverted screen test, LDN-212854-treated animals demonstrated significantly reduced time hanging on the screen compared to controls (Fig. 3F).

**Figure 3.**
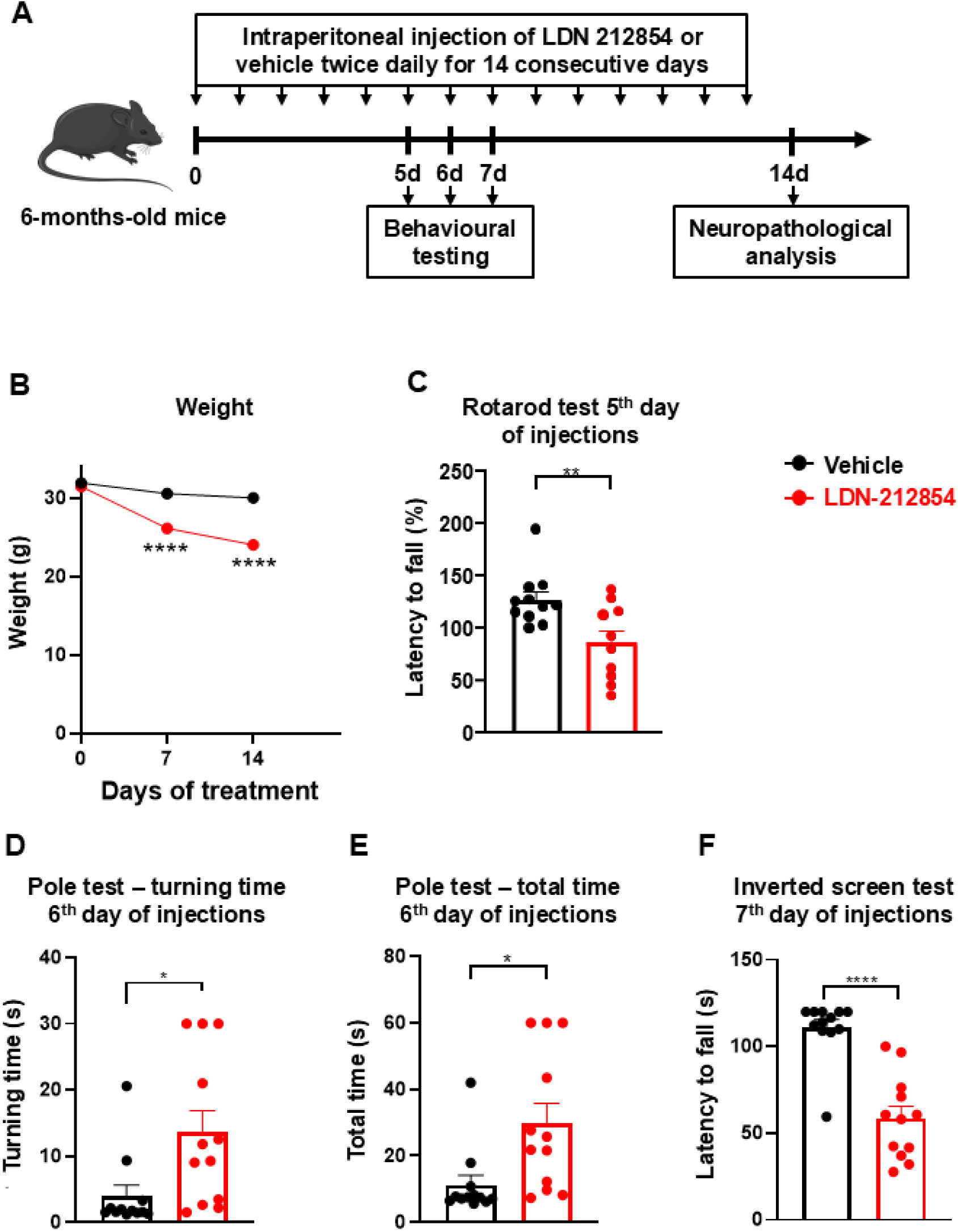
Pharmacological BMP receptor blockade with LDN-212854 induces motor deficits. **(A)** Schematic overview of the experimental design illustrating LDN-212854 or vehicle administration and subsequent behavioral and histological analyses. **(B)** LDN-212854 leads to a significant weight loss **(C)** Motor coordination assessed by the accelerating rotarod test during treatment, was significantly reduced by LDN treatment compared to vehicle-treatment. **(D, E)** The pole test performance parameters **(D)** turning time (latency to orient downward) and **(E)** total time to descend the pole was increased in LDN treated mice. **(F)** Muscle strength as measured as measured by the latency to fall in the inverted screen test was reduced in LDN treated mice. Statistical analysis was conducted using unpaired t-tests. Significance levels are indicated as follows: *p < 0.05; **p < 0.01; ****p < 0.0001.

Neuropathological analyses revealed deficits in both the striatum and the SNpc of LDN-212854-treated mice. Striatal TH fluorescence intensity was significantly reduced in these animals, indicating dopaminergic terminal loss or dysfunction (Fig. 4A, B). While in the SNpc DA neuronal number as measured by NeuN/TH double stained cells was not changed between treatment groups, the TH^+^ area was reduced by LDN-212854 treatment (Fig. 4D, E). Total p129-α-synuclein area was significantly increased in the SNpc as well as the percentage of TH+ area containing p129-α-synuclein expression (Fig. 4F, G).

**Figure 4.**
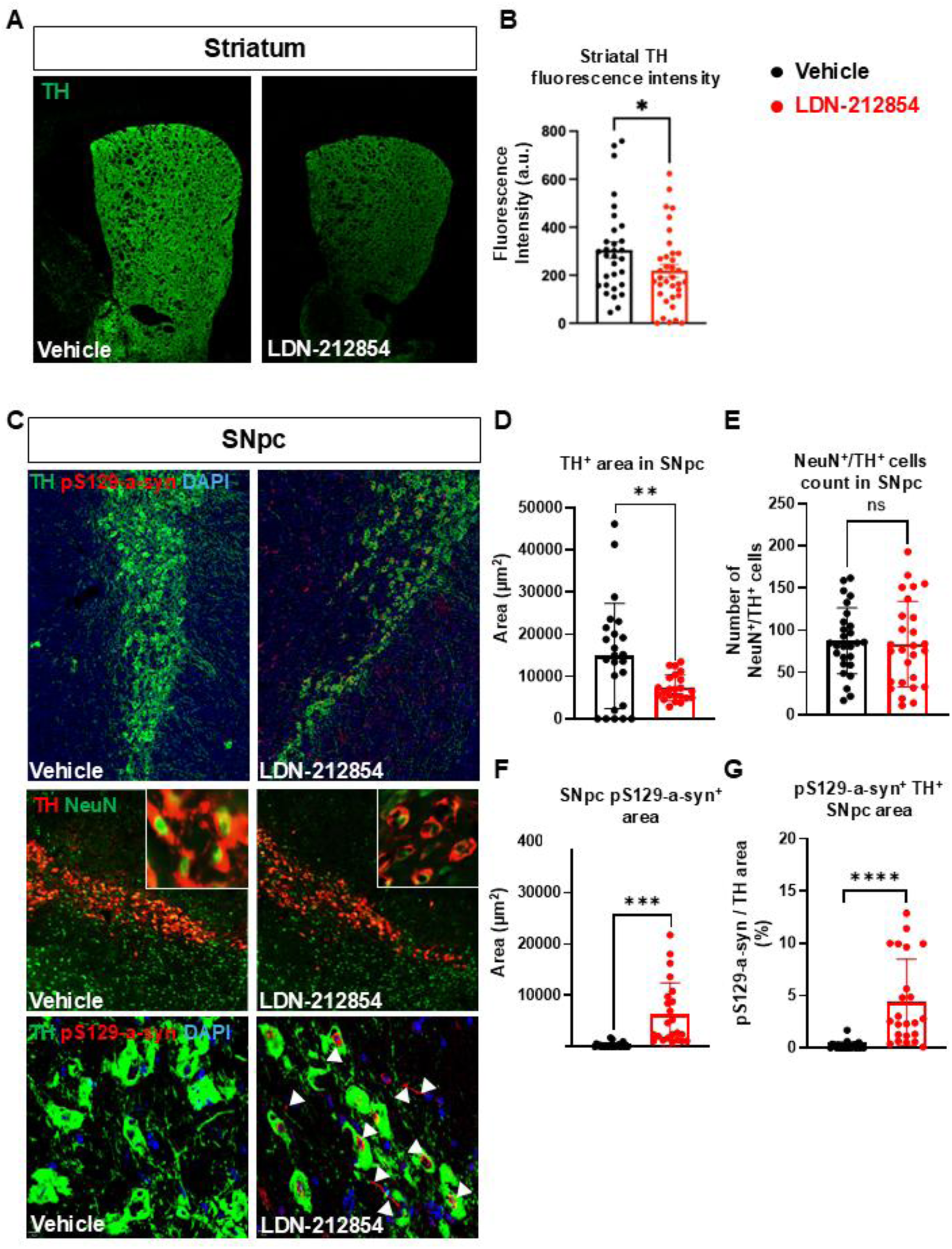
Pharmacological BMP receptor blockade with LDN-212854 causes TH and p129-α-synuclein pathological alterations. (A, **C)** Immunostaining of TH and p129-α-synuclein in LDN-212854 and vehicle treated animals. **(A, B)** Relative optical density of striatal TH signal, is significantly reduced in the treatment group. **(C)** While the area of TH staining was significantly reduced by the treatment, the number of NeuN/TH positive cells was not **(G)**. **(E, F)** p129-α-synucelin area are significanatly increased in the SNpc and in the TH+ area. Statistical analysis was conducted using unpaired t-tests. Significance levels are indicated as follows: *p < 0.05; **p < 0.01; ***p < 0.001.

Taken together, our pharmacological experiments extend our genetic findings suggesting that BMP signaling has also in adulthood an essential role in preserving normal motor behaviour and preventing DA neurons α-synuclein pathology.

### Concomitant delivery of BMP5/7 with α-synuclein PFFs confers neuroprotection against motor deficits and neuropathological abnormalities

We investigated the neuroprotective effects of BMP5/7 against α-synuclein PFFs and compared BMP5/7’s effects to those of GDNF treatment. While it has been previously reported that GDNF exhibits neuroprotective effects when simultaneously injected with α-synuclein PFFs its potential to prevent PFFs-induced motor abnormalities is unknown ^36^. We performed unilateral stereotactic injections in C57BL/6JRccHsd mice. Mice, aged 5 months at the time of injection, were chosen because α-synuclein PFFs injections in younger animals did not produce robust pathology in our hands (data not shown). α-synuclein PFFs (5 µg per dose) were injected into the striatum, while AAV1/2 viral vectors expressing BMP5/7 or GDNF were injected into the SNpc. As a control group, we utilized mice that received unilateral intrastriatal injections of α-synuclein monomers (5 µg per dose). Motor function and neuropathological changes were evaluated for four months post-injections (Fig. 5A, B).

**Figure 5.**
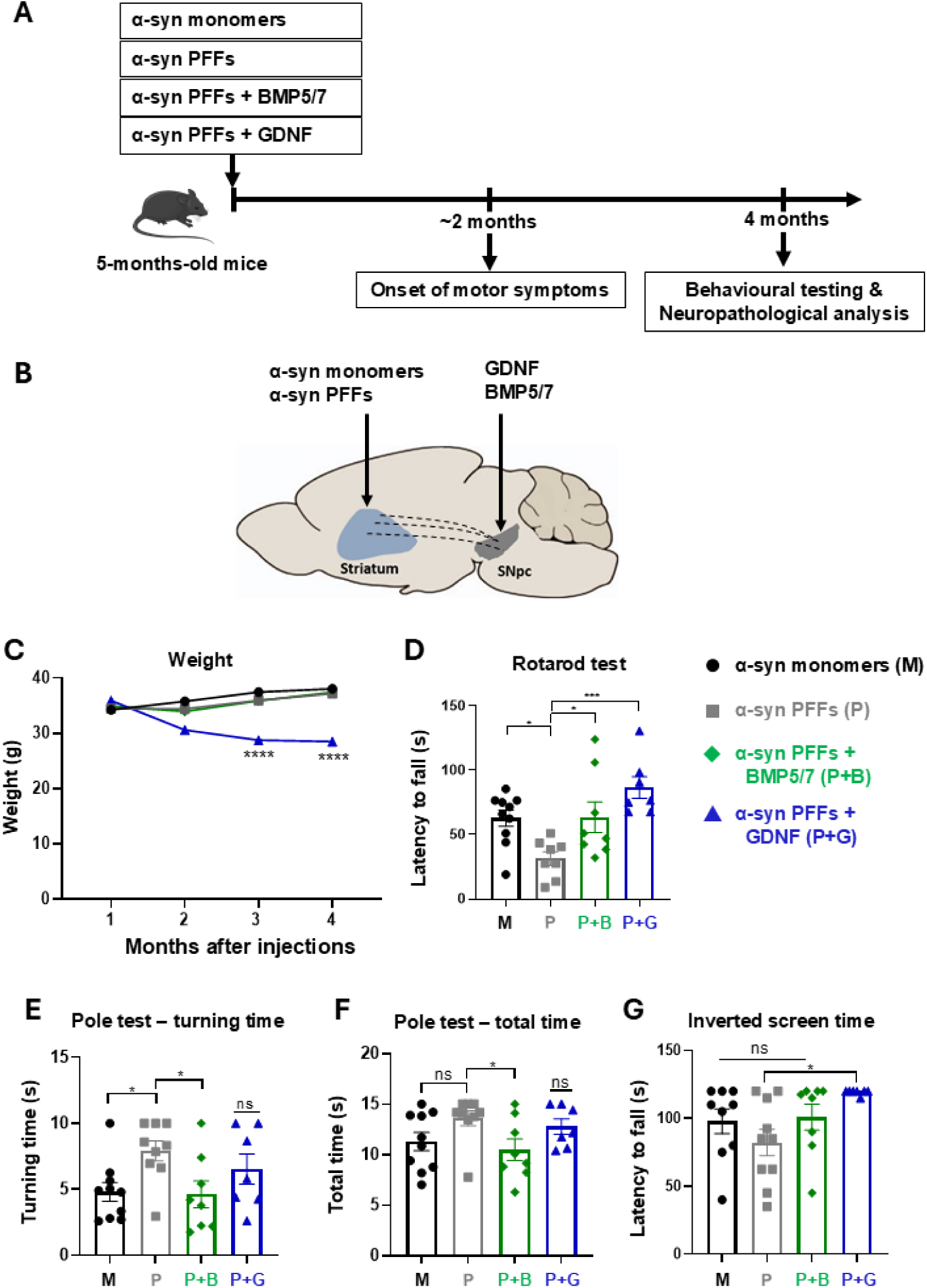
BMP5/7 and GDNF treatments prevent a-synuclein PFF-induced motor deficits. **(A)** Experimental timeline and **(B)** Stereotactic injection sites. **(C)** α-synuclein-PFFs applied together with GDNF leads to reduced body weight. **(D)** In the rotarod test, BMP5/7 and GDNF applied together with α-synuclein PFFs reduced latency time to fall compared to animals treated with α-synuclein PFFs alone. **(E, F)** In the pole test BMP5/7 significantly reduced the increase in turning and total time caused by α-synuclein-PFFs treatment. **(G)** In the inverted screen test, GDNF increased significantly the latency to fall induced by α-synuclein PFFs. Data are presented as mean ± SEM, with each point representing an individual animal. Statistical significance was evaluated using one-way ANOVA followed by Dunnett’s post-hoc test, comparing all groups to the PFF-injected control. Significance thresholds are denoted as follows: *p < 0.05, ***p < 0.0005, ****p < 0.00005, ns – not significant.

Measuring the animals’ weights revealed that GDNF treatment led to a significant reduction in weight (Fig. 5C), as previously reported in humans ^37, 38^ and rats ^39^. In contrast, animals in all other groups maintained stable body weight post-injections (Fig. 5C). In the rotarod test, α-synuclein PFFs exposure resulted in a reduced latency to fall, which was significantly improved by both BMP5/7 and GDNF (Fig. 5D). In the pole test, BMP5/7 significantly improved both turning time and total descent time, whereas GDNF’s effects did not reach statistical significance (Fig. 5E, F). In the inverted screen test, while PFFs injections did not result in a significant decline in latency compared to monomers, GDNF-treated mice exhibited a significantly higher latency to fall than those receiving PFFs alone (Fig. 5G).

Subsequently, we investigated the cellular underpinnings of the neuroprotective effects of the tested neurotrophic factors. Striatal mean fluorescence intensity of TH immunostaining, used as a readout for the intactness of nigrostriatal projections, was significantly reduced by α-synuclein PFFs compared to the control group. This effect was significantly prevented by both BMP5/7 and GDNF (Fig. 6A, C). While in the SNpc the α-synuclein PFFs treatment group showed a significant decrease in TH expression area, co-administration of both BMP5/7- and GDNF-prevented this significant loss (Fig. 6A, D). DA neuronal numbers as quantified by NeuN/TH double positive cells did not show any significant differences between the treatment groups (Fig. 6 E). Both BMP5/7 and GDNF-treated groups showed a significant reduction in the total area of pS129-α-synuclein in the SN (Fig. 6F) and in the pS129-α-synuclein area within the TH^+^ staining (Fig. 6G) compared to α-synuclein PFFs-injected mice.

**Figure 6.**
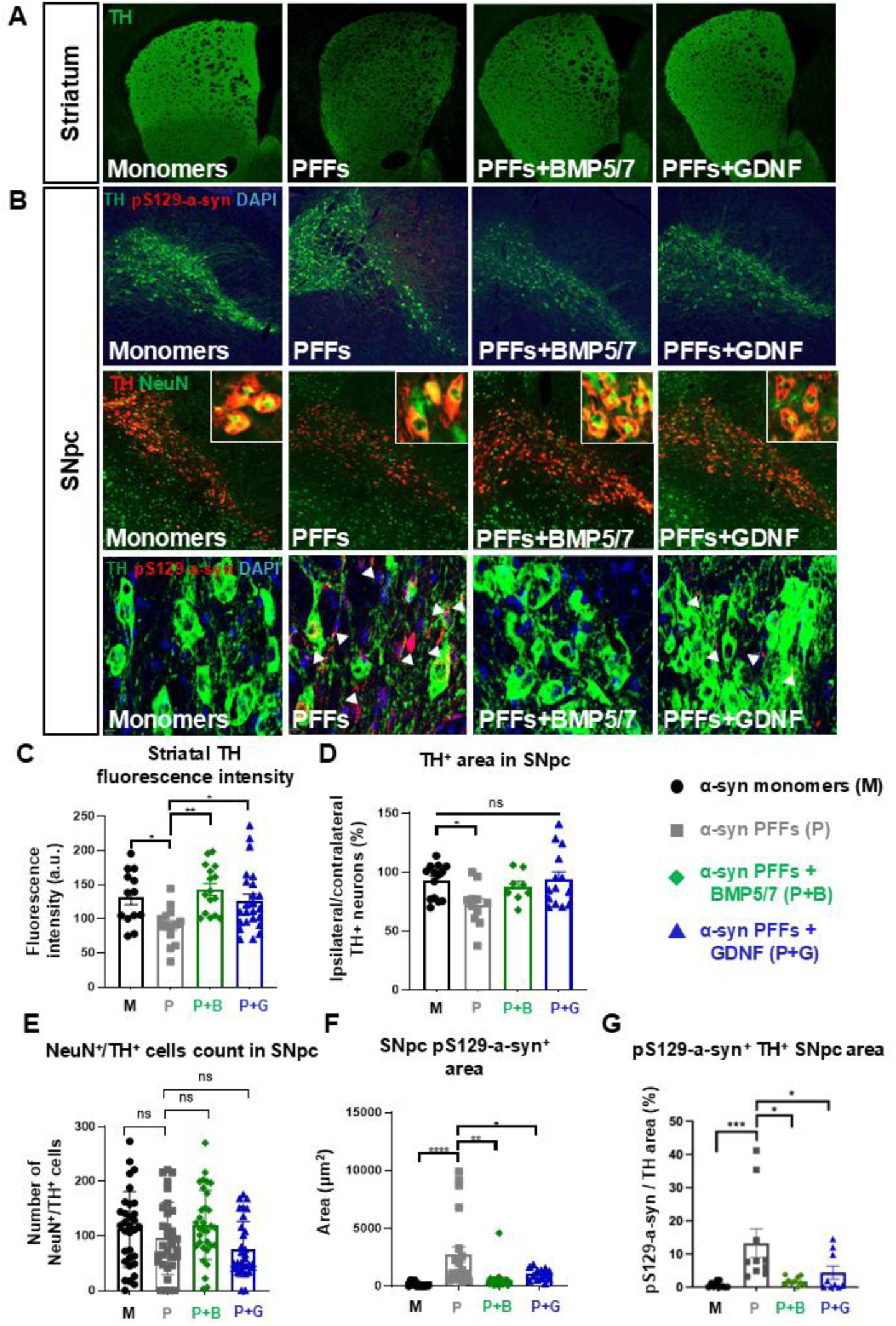
BMP5/7 and GDNF treatments protect against α-synuclein-PFFs-induced pathology. (A, **B)** Immunostainings of striatum and SNpc. **(C)** BMP5/7 and GDNF prevent α-synuclein PFFs-induced reduction in striatal TH fluorescence intensity. **(D)** α-synuclein PFFs-caused decrease of TH, expressed as a percentage relative to the contralateral (uninjected) hemisphere, was averted by BMP5/7 and GDNF. **(E)** DA neuronal numbers as quantified by NeuN/TH double staining did not differ between groups. **(F, G)** BMP5/7 as well as GDNF reduced α-synuclein PFFs increases in p129-α-synucelin area in the SNpc and in the TH+ area. All images were acquired post-mortem from 9-month-old mice, 4 months after stereotactic injection of a-syn PFFs and monomers, following behavioral assessments and tissue processing. Statistical analysis was performed using one-way ANOVA followed by Dunnett’s post-hoc test for comparisons to the PFF-injected control group. Statistical significance is indicated as follows: *p < 0.05, **p < 0.01, **** p < 0.0001.

### BMP5/7 delivered after motor symptom onset ameliorates motor dysfunctions and neuropathological damage induced by α-synuclein PFFs

Next, we assessed the therapeutic efficacy of BMP5/7 and GDNF treatment when administered following the onset of motor symptoms which we observed two months after striatal α-synuclein PFF injection (Fig. 7A). As in our previous experiments testing the neuroprotective effects of BMP5/7 (Fig. 5, 6), we performed unilateral stereotactic injections in aged 5 months old C57BL/6JRccHsd mice. α-synuclein PFFs (5 µg per dose) were delivered into the striatum, while AAV1/2 viral vectors expressing BMP5/7 or GDNF were injected into the SNpc at the onset of motor symptoms. Mice that received unilateral intrastriatal injections of α-synuclein monomers (5 µg per dose) served as a control group.

**Figure 7.**
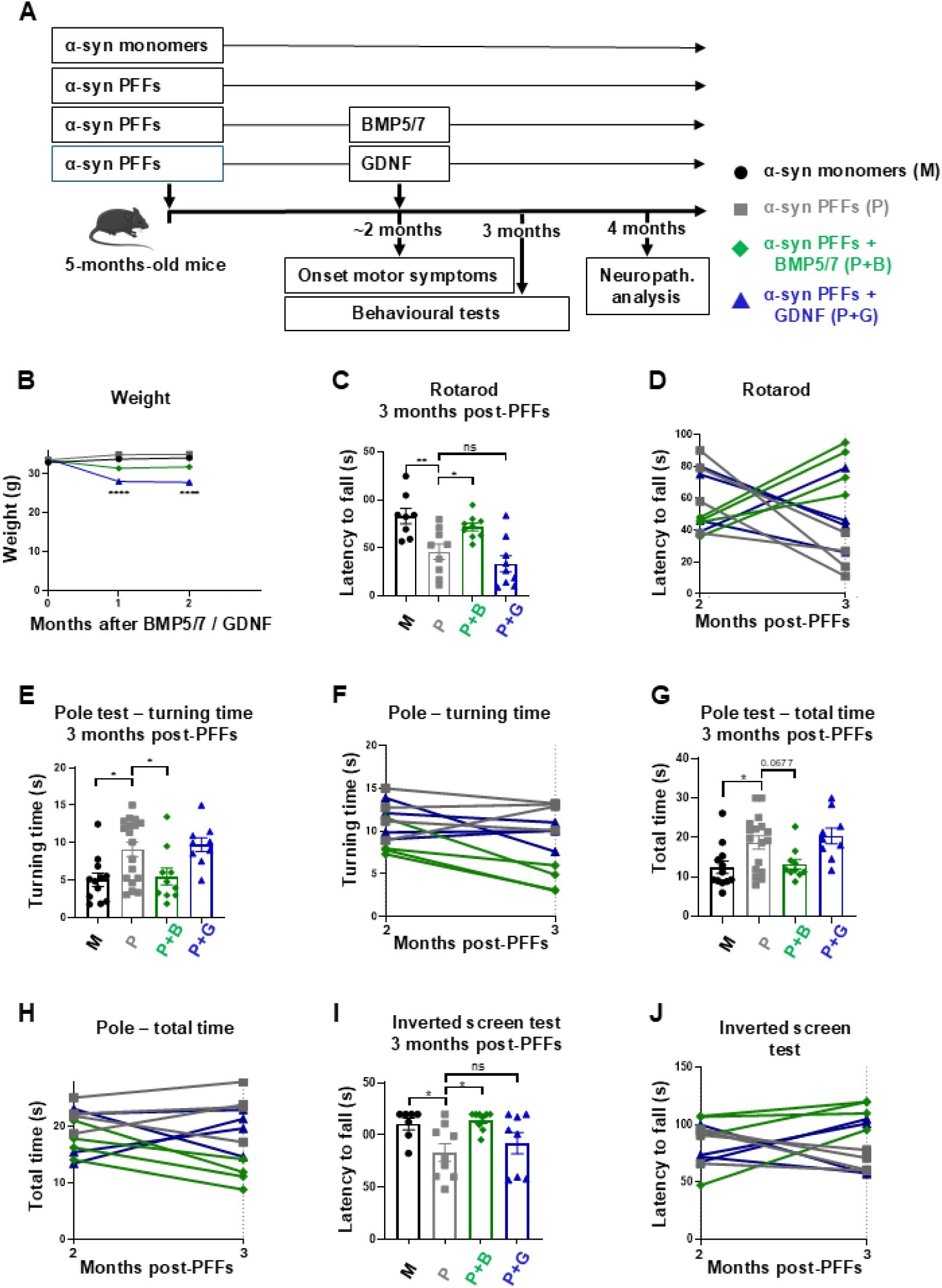
BMP5/7 treatment rescues motor deficits when applied after the onset of a-synuclein PFFs-induced motor symptoms. **(A)** Experimental design **(B)** GDNF treated animals show significant weight loss over time. BMP5/7 effects measured in the **(C)** Rotarod test **(E)** pole test turning time and **(I)** inverted screen test indicate that BMP5/7 significantly reduces behavioral defects induced by α-synuclein PFFs. **(G)** BMP5/7 effects measured by the total descent time in the pole test, showed a trend, but did not reach statistical significance. **(D, F, H, J)** depict the performance of individual mice for each test. **(C-J)** GDNF treatment did not reverse α-synuclein PFFs induced motor abnormalities in the performed tests. Statistical analysis was conducted using one-way ANOVA followed by Dunnett’s post-hoc test to compare all treatment groups to the PFF-injected control group. Significance levels are indicated as: *p < 0.05, **p < 0.01, ***p < 0.001.

Consistent with our previous experiments (Fig. 5C), GDNF treatment led to a reduction in body weight (Fig. 7B). Two months after α-synuclein PFF injection, mice exhibited clear motor impairments, including reduced latency to fall in the rotarod test (Supplementary Fig. 1A), prolonged turning time and total descent time in the pole test (Supplementary Fig. 1B, C), and reduced latency to fall in the inverted screen test (Supplementary Fig. 1D). One month following neurotrophic factors treatment, in the rotarod test, BMP5/7-treated mice demonstrated significantly increased latency to fall compared to α-synuclein PFF-injected mice (Fig. 7C, D). In the pole test, BMP5/7 treatment reduced significantly both turning time (Fig. 7E, F) and total descent time (Fig. 7G, H) to levels closer to control mice (Fig. 7E-H). In the inverted screen test, BMP5/7-treated mice demonstrated significantly improved latency to fall (Fig. 7I, J). Unlike the BMP5/7 group, the GDNF treatment group showed no statistically significant differences compared to the α-synuclein PFFs control group across any of the tests.

To elucidate the cellular mechanisms associated with the therapeutic effects of BMP5/7, we assessed neuropathological changes by immunostaining. First, we confirmed that viral expression of BMP5/7 and GDNF are indeed localized in the SNpc following stereotactic injection (Supplementary Fig. 2). Striatal TH did not show any significant differences between treatment groups (Fig. 8C). However, α-synuclein PFFs injections led to a marked increase of striatal p129-α-synuclein, which was not observed in the BMP5/7 treatment group (Fig. 8D). In the SNpc, the TH+ area was decreased in the group exposed to α-synuclein PFFs, but not in the group treated with BMP5/7 (Fig. 8E). While α-synuclein PFFs led to a decrease in the number of DA neurons as visualized by NeuN/TH double positive staining, the BMP5/7 treatment group did not show this reduction (Fig. 8F). α-synuclein PFFs induced a significant increase in the total area of p129-α-synuclein in the SNpc (Fig. 8G) as well as an increase in the percentage of p129-α-synuclein expression area within the TH expression domain (Fig. 8H). In contrast, in the BMP5/7 treatment group there was no increase in p129-α-synuclein expression (Fig. 8G, H). In contrast to the BMP5/7 treatment group, the GDNF treatment group did not show any statistically significant changes compared to the α-synuclein PFFs group (Fig. 8A-G).

**Figure 8.**
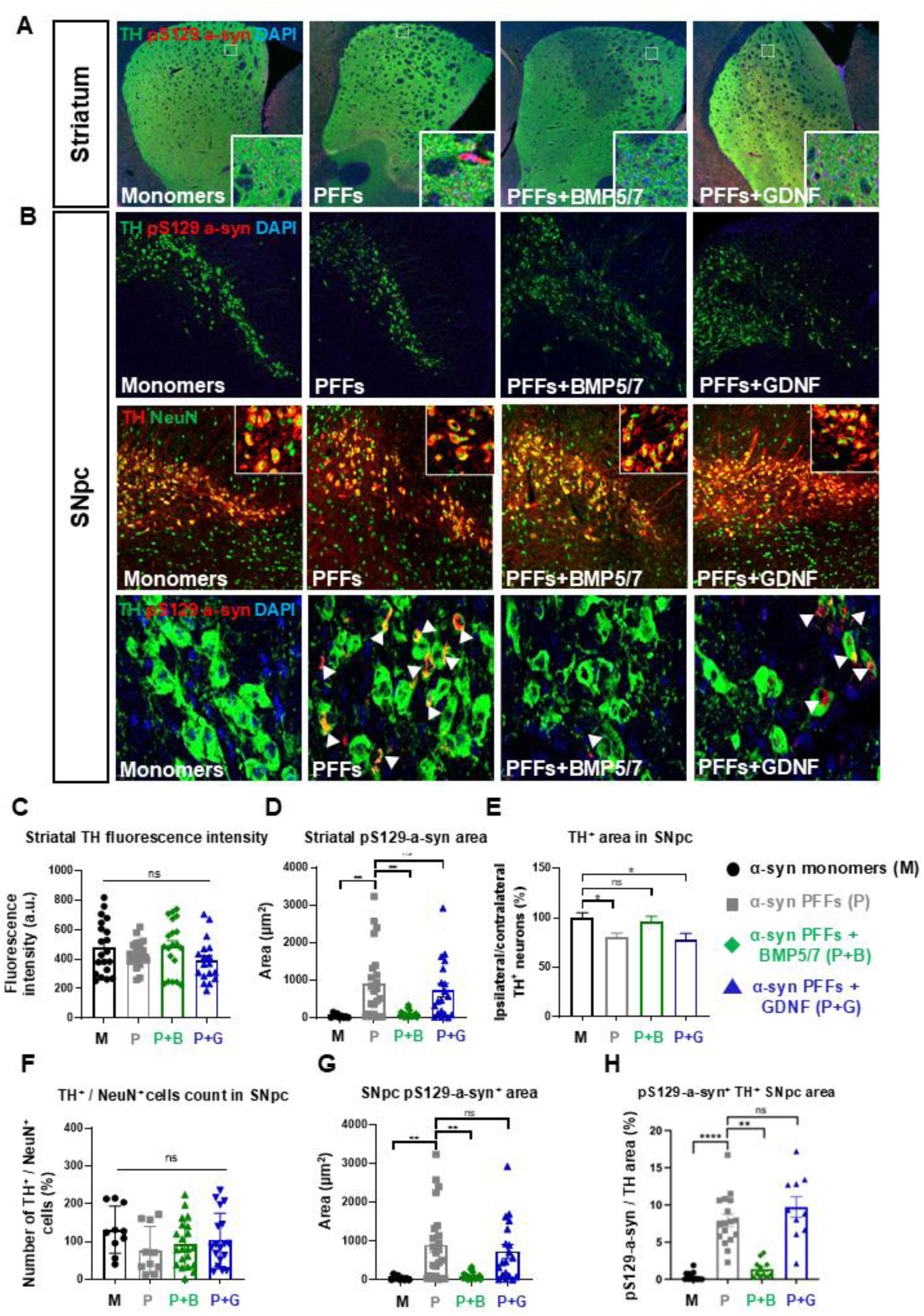
BMP5/7 treatment show neurorestorative effects when applied after the onset of a-synuclein PFFs-induced motor deficits. (A, **B)** Representative immunofluorescence images of the striatum and SNpc. **(C)** Striatal TH staining intensity did not show significant differences between the treatment groups. **(D)** BMP5/7 treatment group significantly reduced striatal p129-α-synuclein area. **(E)** BMP5/7 treated animal do not show α-synuclein-PFFs induced reduction in the SNpc TH area. **(F)** BMPP5/7 treatment normalizes the number of SNpc TH^+^ neurons as a percentage relative to the contralateral hemisphere. **(G)** BMP5/7 treatment group does not show an increase in the p129-α-synuclein total area and **(H)** the percentage of area colocalizing with TH. GDNF treatment group did not show a significant difference to the α-synuclein PFFs treatment group in all measured parameters **(C-H)**. All measurements were obtained four months following intrastriatal PFF injection, with BMP5/7 or GDNF administered two months post-PFFs, after motor symptoms had emerged. Statistical analysis was conducted using one-way ANOVA followed by Dunnett’s post-hoc test. Significance levels are indicated as follows: ns – not significant, *p < 0.05, **p < 0.01, ***p < 0.001, ****p < 0.0001.

We conclude that BMP5/7 treatment at the onset of α-synuclein induced motor impairments show therapeutic effects on PD-associated motor abnormalities and neuropathology.

## Discussion

Here we report that the canonical BMP pathway is involved in the risk of PD. The biological significance of the human data was validated by results showing that the genetic and pharmacological blockage of the canonical BMP signaling leads to PD-associated motor impairments and neuropathological changes. Moreover, we demonstrate the neuroprotective effects of BMP5/7 when applied together with α-synuclein PFFs. Finally, we also provide evidence of the neurorestorative effects of BMP5/7 applied at the onset of motor symptoms, two months after PFFs exposure.

In this study, we systematically investigated the role of the BMP pathway in PD using complementary genetic approaches, including single-variant association analyses, pathway-specific polygenic risk scoring, and rare variant burden testing. Our results provide for the first time converging evidence that genetic variation within the BMP pathway contributes to PD susceptibility primarily at the polygenic level.

Although several common variants showed nominal associations with PD risk and AAO, none survived multiple testing correction, indicating that individual BMP pathway variants are unlikely to exert strong independent effects. In contrast, pathway-level PRS analyses revealed a robust and reproducible association with PD risk, which remained significant in sensitivity analyses excluding genes encoding extracellular antagonists or the intracellular antagonists SMURFs, given their known pleiotropic roles across multiple signaling pathways. These findings suggest that the cumulative effect of many small-effect variants within the BMP pathway contributes meaningfully to disease susceptibility. The significant association between BMP pathway PRS and AAO observed in the dataset including proxy cases was modest and attenuated after removing genes encoding extracellular antagonists. This further suggests that while the BMP pathway may influence disease risk, its role in modulating disease onset may be limited. Our rare variant analyses provided limited evidence for a contribution of low-frequency variants. The lack of consistent replication across datasets suggests that larger sample sizes may be required to clarify the role of rare variants in this pathway.

Our results indicate that the loss of BMP/SMAD signaling in DA neurons leads to motor impairments associated with p129 α-synuclein accumulation. We previously demonstrated that genetic inactivation of SMAD1, an essential mediator of intracellular canonical BMP signaling, during mouse embryogenesis results in α-synuclein accumulation, aggregation, and the loss of DA neurons in adulthood ^25^. Here, we extend our knowledge by demonstrating that a loss of SMAD1 leads to motor impairments, and we shed light on the cellular mechanisms underpinning the protective effects of the BMP/SMAD signaling against PD-associated pathology. In our previous study ^25^, we inactivated SMAD1 in both neurons and glial cells as we used a Nestin-Cre driver ^40^. Consequently, those experiments did not resolve whether BMP/SMAD signaling activation in neurons or in glia protects against α-synuclein accumulation. Glia cells are well known to be responsive to BMPs. Therefore, an indirect mechanism, by which BMPs activate the canonical SMAD signaling cascade in glia, which then indirectly protects neurons against α-synuclein-associated neurodegeneration would be conceivable ^41^. By employing a DA neuron-specific Cre-driver line, we reveal an essential role of SMAD1 in safeguarding DA neurons against α-synuclein pathology and motor function and that this process is cell-autonomous and not mediated indirectly via glia cells.

Our pharmacological BMP pathway manipulation indicates that BMP/SMAD blockage in adult mice leads to PD-associated pathology. The strength of using the DAT-Cre driver is that it enabled us to identify cell autonomous functions of SMAD1 in DA neurons. However, a limitation of this approach is that the long period between embryonic genetic manipulation and adult phenotype analysis allows for the development of compensatory mechanisms, which complicates the interpretation. Our pharmacological experiments clearly demonstrate a role of BMP/SMAD signaling in adulthood for safeguarding DA neurons against PD-associated pathology and motor abnormalities. By utilizing acute pharmacological blockage, we bypassed these developmental confounds and demonstrate that the BMP/SMAD pathway must remain active in adulthood to safeguard DA neurons against PD-associated pathology and subsequent motor abnormalities. Finally, our genetic and pharmacological experiments suggest that the direct modulation of α-synuclein pathology plays an essential role in the neuroprotective and neurorestorative effects of BMP5/7.

Our results indicate that BMP5/7 protects DA neurons against α-synuclein PFFs-induced toxicity *in vivo.* Previous studies have shown that BMPs show beneficial effects in toxin-based *in vitro* and *in vivo* PD models ^23^. Although toxin-based PD models are very efficient in inducing acute loss of DA neurons, they do not recapitulate the slow neurodegenerative process associated with α-synuclein pathology, found in familiar or idiopathic PD ^42^. Two independent studies aiming to test the neuroprotective effects of BMPs against α-synuclein induced neurodegeneration *in vivo*, demonstrated the effectiveness of BMPs in preventing neuropathological changes and motor abnormalities. BMPs were efficient when administered simultaneously with virally overexpressed wild-type or mutated α-synuclein in mice or rats ^24, 25^. The α-synuclein viral overexpression PD model, is particularly relevant to recapitulate pathophysiological processes found in patients with α-synuclein gene duplications/triplications or α-synuclein mutations ^43^. However, the supraphysiological α-synuclein levels induced in the viral overexpression paradigm limits its validity for idiopathic PD. To address these limitations the α-synuclein PFFs PD model was developed. Here, injected α-synuclein PFFs, acting as seeds, are used to induce endogenous α-synuclein aggregation that serve as further templates within the host neurons, reflecting the process thought to drive pathology in sporadic PD. It mimics the “prion-like” spreading of α-synuclein pathology through interconnected brain regions, believed to occur in idiopathic PD and is therefore regarded as a prime model for idiopathic PD ^20, 42, 43^. Thus, the efficacy of BMP5/7 in the α-synuclein PFFs-based PD *in vivo* model, demonstrates their therapeutic potential for idiopathic PD.

A major finding of our study is that BMP5/7 showed neurorestorative effects when applied *after* the onset of motor symptoms caused by already damaged dopaminergic neurons. At the onset of the PD symptoms, there is about 80% decrease in dopamine content in the striatum but still about 50% of nigral DA cells are viable. The fact that these neurons have not undergone cell death yet, opens a critical time window at the time of diagnosis for neurorestoration until these neurons die during the disease. However, neuropathological changes are already widely spread at the time of PD diagnosis ^44, 45^. Therefore, testing the neurorestorative effects of drug candidates in preclinical models in which neurons are already undergoing neuropathological changes is essential to assess their clinical potential. Whereas in preclinical experiments, the efficacy of neurotrophic factors to protect healthy neurons against the development of α-synuclein-induced toxicity has been well documented, their therapeutic effect in degenerating neurons has been largely unknown ^16^. Therefore, our findings that BMP5/7 can robustly reverse neuropathological changes as well as motor abnormalities after α-synuclein PFFs induced pathology provide yet, the best available indication for the disease-modifying potential of these neurotrophic factors in PD.

While BMP5/7 showed neurorestorative effects in our experiments, GDNF did not. A recent clinical 1b trial has presented further safety of GDNF and promising results for its efficacy, that motivated further clinical testing ^46^. With hopefully positive results in further clinical trials, GDNF will show the way as the first neurotrophic factor efficient in PD. However, because previous clinical trials testing GDNF protein family members failed to meet their endpoints, concerns regarding GDNF efficacy persist, highlighting the need for further stringent preclinical validation ^16^. GDNF has shown robust neuroprotective and neurorestorative effects in preclinical toxin-based *in vivo* PD models ^15–17^. However, reports about its efficacy in α-synuclein-based *in vivo* PD models are mixed. While GDNF showed *in vivo* neuroprotective effects against α-synuclein PFFs ^36^, it did not protect against viral overexpression of α-synuclein ^47, 48^. Our results of the neuroprotective effects of GDNF when applied *together* with α-synuclein PFFs are in accordance with previous results ^36^. We extend these findings by demonstrating that GDNF protects against α-synuclein PFFs-induced motor abnormalities. However, GDNF did not show any neurorestorative effects when compared head-to-head to BMP5/7. We cannot exclude that when using a different dosage of GDNF we would be able demonstrate its neurorestorative effects. However, the GDNF dosage we used for the delayed application was efficient when administered simultaneously with α-synuclein PFFs. Also, did we observe sufficient coverage of the SNc with GDNF. Consistent with our results GDNF did not show any neurorestorative effects in a proteasome inhibition mouse *in vivo* PD model. In these experiments different genetic approaches to upregulate endogenous GDNF and AAV5-GDNF did not restore neural damage or behavioural abnormalities ^49^. While the focus of our current study was on BMPs, the direct comparison with GDNF indicates that a further analysis of the neurorestorative effects of GDNF is warranted.

Taken together, we conclude that impairments in the BMP/SMAD signalling pathway lead to α-synuclein associated neuropathological changes and motor abnormalities and increase the risk of PD. Moreover, our findings indicate that BMP5/7 show therapeutic effects in neurons affected by α-synuclein-associated pathology. Thus, our findings indicate BMPs are promising disease-modifying drug candidates for PD.

## Material and Methods

### Genetic Data and Sample Quality Control

For genotyping data, samples from the GP2 cohort were generated as part of the Global Parkinson’s Genetics Program (GP2; release 10). Quality control (QC) procedures for both samples and variants have been previously described (https://github.com/GP2code/GenoTools). The UKB genetic dataset comprises 487,409 samples, which were phased and imputed using the Haplotype Reference Consortium and the combined UK10K + 1000 Genomes reference panels ^50^. Only unrelated individuals of European ancestry were retained for downstream analyses. To merge the GP2 and UKB datasets, high-quality imputed variants (R² > 0.8) shared between both cohorts were selected. Within each cohort, variants were independently filtered based on minor allele frequency (MAF > 0.01), genotype missingness (GENO < 0.05), and Hardy–Weinberg equilibrium in controls (P > 5 × 10⁻⁶). After merging, these QC filters were reapplied to ensure the integrity and consistency of the combined dataset. The final dataset included 3,597 cases and 35,970 controls in analyses without proxy cases, and 18,611 cases and 186,110 controls when proxy cases were included.

WGS data from the AMP-PD Study Group (release v4; https://amp-pd.org/) underwent initial QC as previously described (https://amp-pd.org/whole-genome-data). Additional filtering steps were applied, including removal of variants with depth of coverage (DP ≤ 25), exclusion of samples with >5% missing genotypes, individuals with excess heterozygosity (F < −0.15 or F > 0.15), and samples with call rate <95%. Individuals of non-European ancestry and those related closer than third-degree (pi-hat > 0.125) were excluded. After QC, the AMP-PD dataset included 3,051 cases and 3,667 controls. UKB WGS samples were processed using the same QC procedures, with the additional requirement of genotype quality (GQ ≥ 25). The resulting dataset included 3,173 cases and 31,722 controls in analyses without proxy cases, and 18,110 cases and 180,962 controls when proxy cases were included.

### BMP Pathway Definition

In this analysis, we focus on the canonical BMP signaling pathway, in which BMP ligands engage type I and II serine/threonine kinase receptors to activate SMAD1/5/8 proteins ^21, 51^.

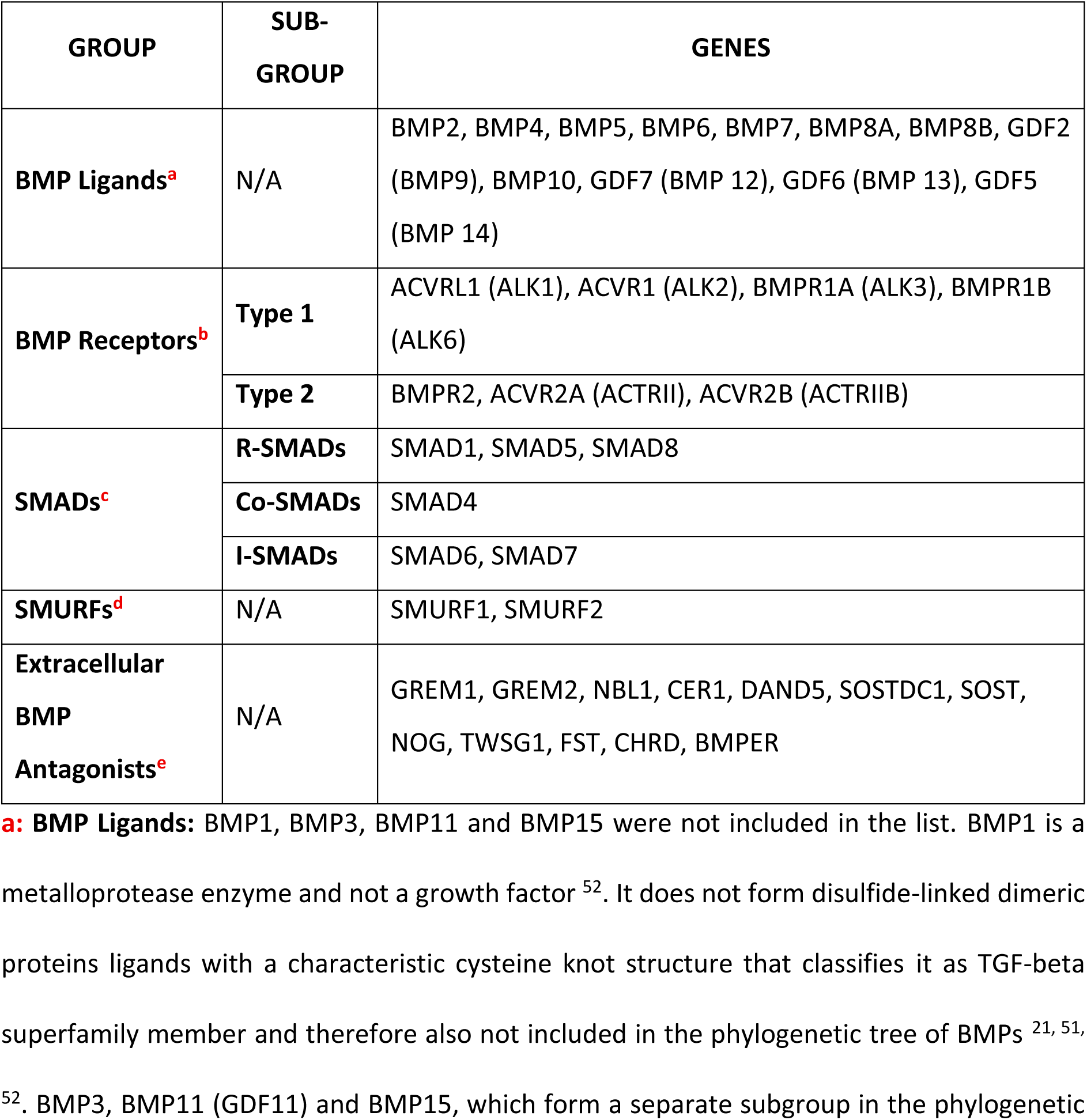
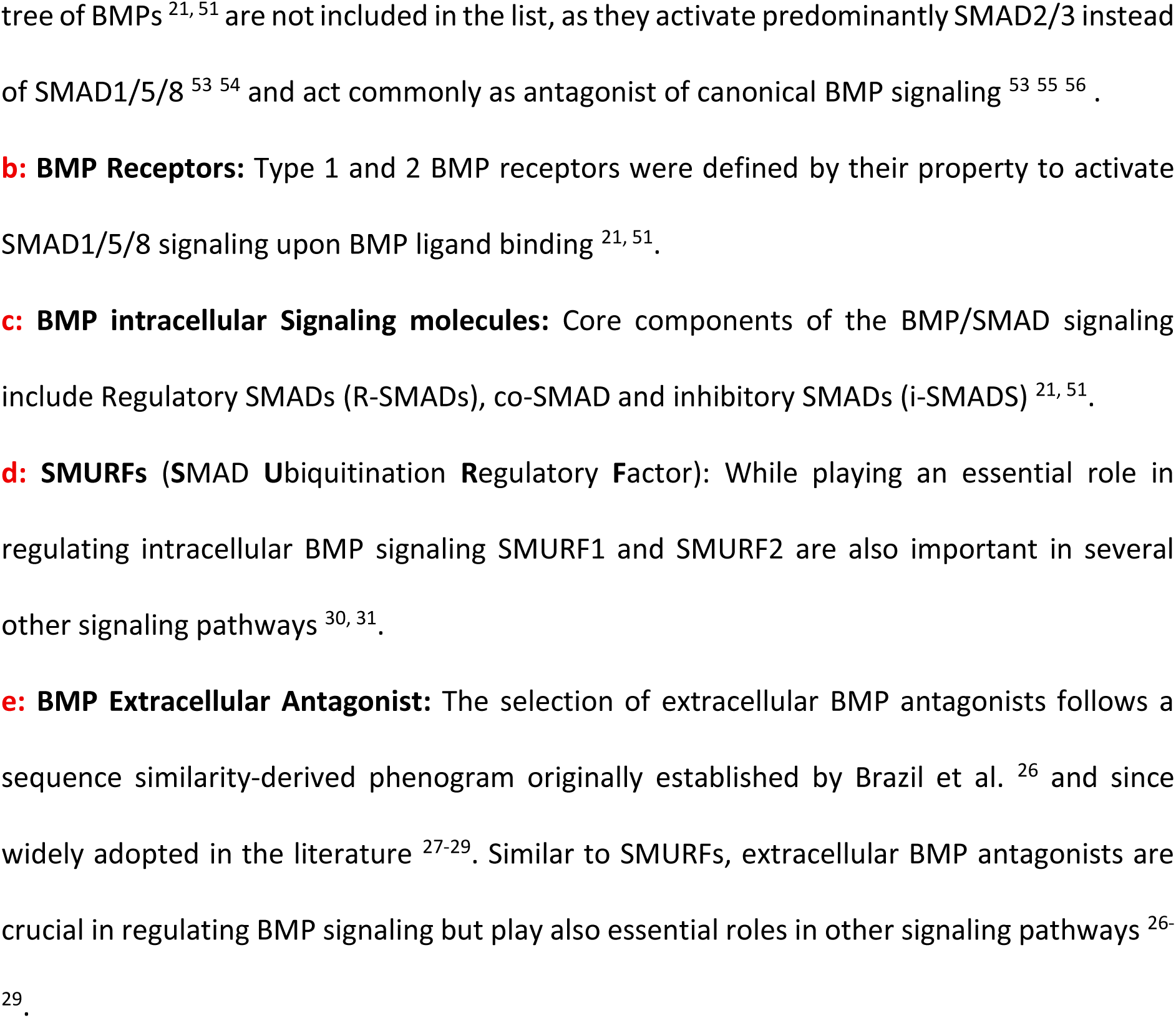

### Genetic Analysis

To assess the association between common variants (MAF > 0.01) in the BMP pathway and PD, genotyping data for BMP pathway genes were extracted and pruned using linkage disequilibrium (LD) parameters of 50 kb windows, a step size of 5 variants, and an r² threshold of 0.2. Logistic regression was performed to evaluate PD risk, adjusting for age, sex, and principal components (PCs). For AAO, linear regression was conducted with adjustment for sex and PCs. All analyses were performed using PLINK v1.9 ^57^.

Pathway-specific PRS were computed using PRSice-2 ^58^, based on the most recent PD GWAS summary statistics ^13, 59^. Standard QC was applied to the summary statistics before PRS construction, including exclusion of variants with MAF < 1%. LD clumping was performed using an r² threshold of 0.1 within a 250 kb window. Regression models were adjusted for age, sex, and the top five PCs. Empirical P-values were generated using 10,000 permutations to assess pathway-level enrichment.

To investigate the contribution of rare variants (MAF < 0.01) within the BMP pathway to PD risk, optimized Sequence Kernel Association Tests (SKAT-O) were performed using the SKAT R package ^60^. SKAT-O evaluates the aggregated effects of rare variants within predefined genomic sets, accounting for both the direction and magnitude of individual variant effects. Gene coordinates were defined using MANE Select transcripts, with an additional 100 base pairs upstream and downstream. Variants were annotated using the Variant Effect Predictor (VEP) ^61^. Burden analyses were conducted across four variants categories: (1) functional variants, defined as nonsynonymous, splice-site, frameshift, and stop-gain variants, (2) loss-of-function variants, including splice-site, frameshift, and stop-gain variants, (3) nonsynonymous variants, and (4) predicted deleterious variants, defined as variants with a Combined Annotation Dependent Depletion (CADD) score >20, representing the top 1% of predicted deleterious variants. The regression model was adjusted for age, sex, and the first five principal components. Meta-analysis across cohorts was conducted using the MetaSKAT package ^62^.

### Animals

*Dat^Cre^* mice have been purchased from Jackson Laboratories (B6.SJL-*Slc6a3^tm1.1(cre)Bkmn^*/J; JAX LAB Strain #:006660) and characterized previously ^34^. *Smad1^fl/fl^* mice have been characterized previously ^32, 33^. *Smad1^fl/fl^* mice were mated to *Dat^Cre^* mice to generate *Smad1^fl/fl^; Dat^Cre^* conditional mutant mice and control littermates.

For the LDN-212854 treatment experiments we used 8-12 weeks old wild-type C57BL/6JRccHsd male animals purchased from Envigo, Israel. For the α-synuclein PFFs treatment experiments we used 5-month-old, C57BL/6RccHsd male mice. All mice were housed in a temperature-controlled (21–23°C) environment, under a 12-h light/dark cycle and had free access to food and water in a pathogen-free animal facility.

### Study approval

All procedures and experimental protocols conducted on the animals were approved by the Institutional Animal Care and Ethics committee at Ben-Gurion University of the Negev (Permit Number: IL-22-04-21-C).

### Genotyping of mouse mutants

Tails were lysed using 200 μL lysis buffer (25mM NaOH, 2mM EDTA) for 30 minutes at 90°C. After cooling down, the reaction was neutralized using 200 μL neutralization buffer (40 mM Tris-HCl). To identify Smad1^fl/fl^ and *Dat^Cre^*mutant mice, a standard multiplex PCR was performed using the primers listed in Table 1. For *Smad1^fl/fl^*, the primers used were S1F400 and LoxP2. For DAT-Cre mutant mice, the primers used were oIMR6625, oIMR6626, and oIMR8292.

**Table 1.**
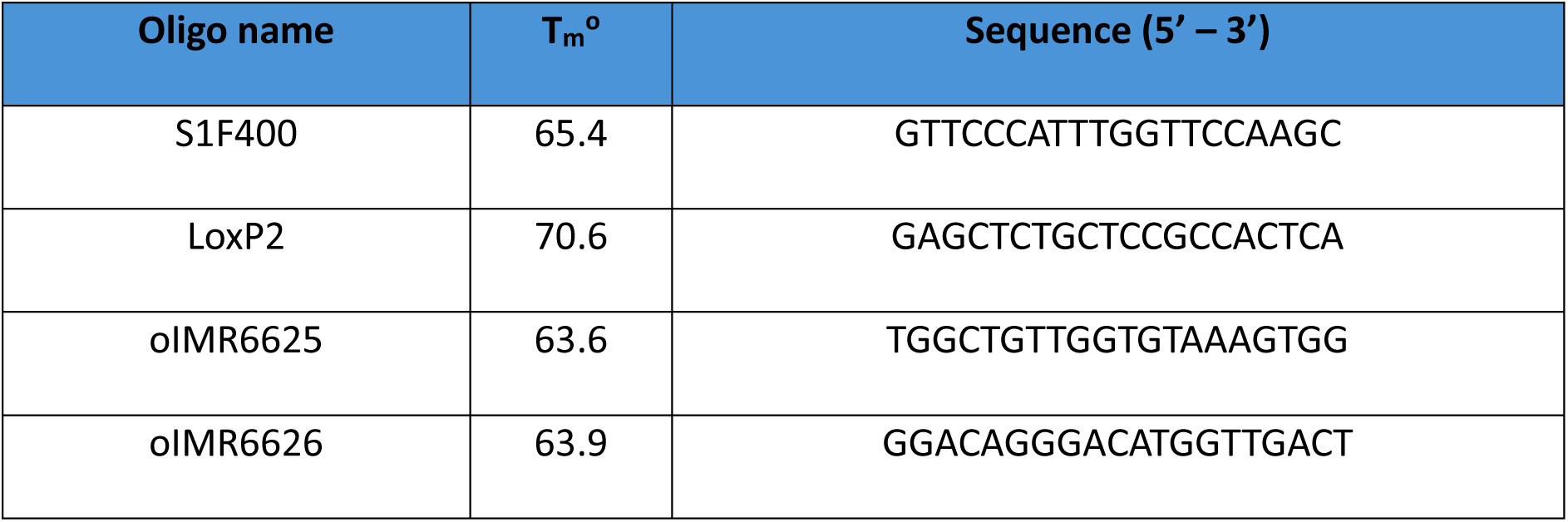
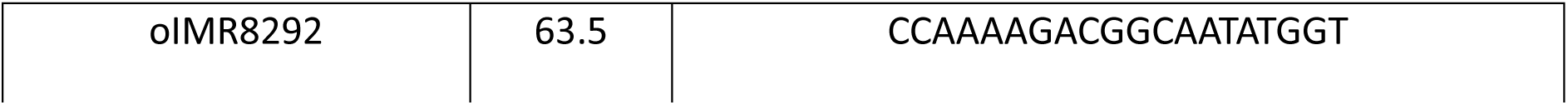
List of primers.

### LDN-212854 treatment

Mice treated with LDN-212854 (MedChemExpress, cat. no. HY-15897) were intraperitoneally injected with 6 mg/kg of LDN-212854 twice a day for two consecutive weeks, with injections given every 12 hours. The control group of mice was injected with a vehicle solution containing 10% DMSO, 40% PEG 300, 5% Tween 80, and 45% saline, at the same times as the LDN-212854-treated group to account for potential effects of the injection and vehicle. The mice were 6 months old at the start of the treatment.

### α-synuclein PFFs preparation and validation

Recombinant mouse α-synuclein monomers (StressMarq) at 5 mg/mL were incubated with glass beads and agitated at 1000 rpm for 7 days at 37 °C to generate PFFs. PFFs formation was validated using a Thioflavin T (ThT) fluorescence assay and transmission electron microscopy (TEM), both before and after sonication, as previously described ^63^. Amyloid fibril formation was assessed using the ThT assay, which detects β-sheet-rich structures through increased fluorescence upon binding. A fresh 1 mM ThT stock solution was prepared in distilled water, filtered through a 0.2 µm syringe filter, and diluted to a final working concentration of 25 µM. α-synuclein monomers and PFFs were thawed on ice immediately prior to use, and samples containing 10 µM PFFs or 100 µM monomers were loaded into black-bottom 96-well plates, sealed, and incubated at 37 °C in a fluorescence microplate reader. ThT fluorescence was measured (excitation: 450 nm; emission: 485 nm) over 72 h. For TEM analysis, 1 µL of each sample was applied to carbon-coated copper grids and incubated for 5 min, followed by staining with 5 µL of 2% (w/v) uranyl acetate for 2 min. Excess stain was removed by blotting, and grids were air-dried for at least 30 min before imaging using a transmission electron microscope. For quantitative analysis, 15 images per sample were randomly acquired across the grid, and fibril length was measured in a blinded manner using ImageJ software.

### Viral vectors

Plasmids were constructed using standard molecular cloning techniques. Briefly, 5 µg of plasmid DNA was linearized by restriction digestion with the appropriate enzymes in 1× CutSmart Buffer (NEB), and digestion efficiency was verified by agarose gel electrophoresis. Linearized plasmids were purified using the NucleoSpin® Gel and PCR Clean-up Kit (Macherey-Nagel) and ligated with the desired inserts using T4 DNA ligase (NEB). Ligated plasmids were transformed into DH5α competent cells, and individual colonies were expanded for plasmid preparation using PureLink™ HiPure Plasmid Midiprep Kit (Invitrogen). Plasmid DNA was dissolved in nuclease-free water, quantified by NanoDrop spectrophotometer, and inserts were confirmed by Sanger sequencing.

AAV1/2 viral vectors were produced in HEK-293T cells using calcium phosphate-mediated transfection as described previously (Groh et al., 2008), with minor adjustments. Seventy-two hours post-transfection, cells were lysed via three freeze-thaw cycles in lysis buffer (150 mM NaCl, 50 mM Tris-HCl), followed by incubation with Benzonase (100 U/mL) for 1 h at 37 °C. Crude lysates were stored at 4 °C for in vitro transduction or further purified for in vivo experiments. Viruses were concentrated and formulated in sterile phosphate-buffered saline (PBS) using Amicon filters (EMD).

### Stereotaxic injections

5-month-old, C57BL/6RccHsd male mice were deeply anesthetized with ketamine/xylazine and placed in a stereotactic head frame (Stoelting, IL, USA). After making a midline incision of the scalp, a burr hole was drilled in the appropriate location for the striatum on the right side of the skull. 2.5 μL of α-synuclein PFFs or monomers were injected at a rate of 0.3 μL/min with a 30-gauge needle on a 25 μL syringe (702 RN, Hamilton Company). The needle was left in place for an additional 5 minutes before being slowly withdrawn from the brain. According to the mouse brain atlas of Paxinos and Franklin, animals were injected at the following coordinates: (AP) +0.4 mm, (ML) −2 mm relative to bregma and (DV) −2.9 mm from the dural surface.

Subsequently, animals were assigned to one of two experimental timelines for intracranial AAV1/2 delivery into the right SNpc at an infusion rate of 0.25 μL/min (total volume: 1.0 μL). For the simultaneous delivery paradigm, animals received the SNpc AAV injections during the same surgical session immediately following the striatal PFF or monomer infusion. For the delayed application paradigm, the AAV injections were administered via a second stereotaxic surgery performed two months post-striatal injection, following the onset of motor symptoms. Depending on the allocated experimental cohort for both paradigms, animals received either AAV1/2-hSyn-hGDNF-V5 alone, or a viral suspension consisting of a 1:1 volume ratio mixture of AAV1/2-hSyn-hBMP5-V5 and AAV1/2-hSyn-hBMP7-V5. The stereotaxic coordinates utilized for the SNpc were AP −3.1 mm, ML −1.4 mm relative to bregma, and DV −4.5 mm from the dural surface. All viral vectors were delivered at the final concentrations and doses specified in Table 2.

**Table 2.**
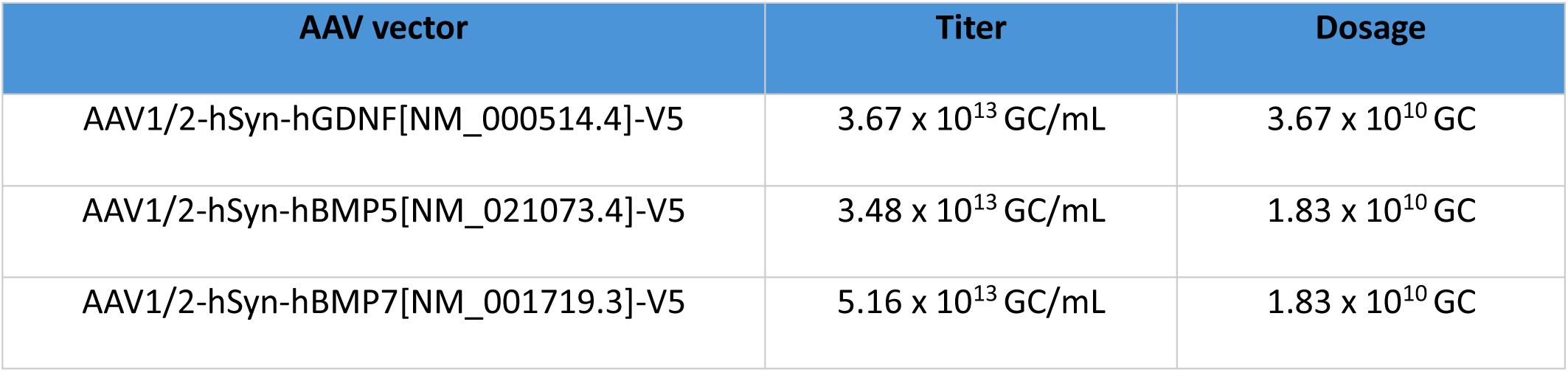
List of AAV vectors used in the study.

### Behavioral analysis

Motor coordination, balance, and strength were evaluated using rotarod, pole, and inverted screen assays. For the rotarod test, mice were placed on an accelerating cylinder (4–40 rpm over 300 s). Latency to fall was recorded automatically via a trip switch and averaged across three trials per animal, with a 15-min inter-trial interval.

Bradykinesia and postural control were assessed using the pole test (60 cm height, 1 cm diameter). Animals were positioned at the apex, facing upward; the time required to turn 180° face-down (T-turn) and the total descent time were recorded. For animals unable to complete the task, default values of 15 s (T-turn) and 30 s (total) were assigned to ensure statistical consistency (Matsuura et al., 1997). Results represent the mean of three trials.

Neuromuscular strength and endurance were quantified via the inverted screen test. Mice were placed on a wire mesh screen, which was inverted 180° over a padded surface. The duration of suspension was recorded for a maximum of 120 s. All behavioral testing was conducted by an investigator blinded to genotype and treatment.

### Tissue processing and immunohistochemistry

Mice were anesthetized with isoflurane via inhalation in a closed chamber according to institutional ethical guidelines and perfused transcardially with PBS followed by 4% paraformaldehyde (PFA). Brains were post-fixed overnight at 4 °C, washed in PBS (3 × 5 min), and cryoprotected in 15% followed by 30% sucrose at 4 °C until sinking. Tissues were embedded in OCT compound (Tissue-Tek 4583), frozen on a thin layer of liquid nitrogen, and stored at –80 °C. Serial 14 µm coronal brain sections were cut on a cryostat and systematically distributed across 6 slide series using a 1-in-6 sampling scheme, with consecutive sections assigned sequentially to each slide in a repeating cycle. Sections were mounted on SuperfrostPlus® slides (Thermo Scientific), air-dried, and stored at –80 °C until use.

For immunohistochemistry, the striatum was represented by 6 slide series generated using a 1-in-6 sampling scheme. The SNpc was represented by 12 slide series, corresponding to two consecutive 1-in-6 sampling cycles (slides 1–6 and 7–12), to ensure full rostrocaudal coverage. Sections were washed in PBS (3 × 5 min) to remove OCT, then briefly fixed in pre-cooled acetone for 10 minutes. Antigen retrieval was performed in 10 mM sodium citrate buffer (pH 6) at 88 °C for 20 min without boiling, then cooled to room temperature. Sections were blocked with 4% normal goat serum (Sigma-Aldrich, Cat. No. G9023) for 1 h at room temperature and incubated overnight at 4 °C with primary antibodies diluted in PBS (see Table 3). The following day, sections were washed (3 × 5 min) and incubated with fluorophore-conjugated secondary antibodies (Alexa 488 and/or Cy3, 1:500, Jackson Laboratories) and DAPI (1:10,000, Sigma Aldrich, Cat. No. D9542) for 1 h at room temperature in the dark. Finally, sections were washed (3 × 5 min) and mounted using Immu-Mount™ (Thermo Scientific, Cat. No. 9990402). All primary and secondary antibodies used are listed in Tables 3 and 4.

**Table 3.** Primary Antibodies.

| Antibody | Host | Working dilution: | Source identifier |
| --- | --- | --- | --- |
| Anti-TH, antibody clone LNC1 | Mouse | 1:500 | MAB318, Millipore |
| Anti-TH antibody | Rabbit | 1:500 | AB152, Millipore |
| Anti-α-synuclein clone 42/α-synuclein | Mouse | 1:500 | 610786, BD Biosciences |
| Anti-pS129 α-synuclein [EP1536Y] | Rabbit | 1:500 | AB51253, Abcam |
| Anti-phosphorylated α-synuclein (pSyn#64) | Mouse | 1:500-1000 | FUJIFILM Wako Pure Chemical Corporation, 015-25191 |
| IBA1 | Rabbit | 1:500 | 019-19741, WAKO |
| GFAP | Rabbit | 1:1000 | Z0334, DAKO |
| LC3b | Rabbit | 1:100 | L7543, Sigma-Aldrich |
| NeuN | Mouse | 1:200 | MAB377x, Chemicon |
| pSMAD1/5/8 | Rabbit | 1:200 | 9511S, Cell Signaling |
| cCaspase-3 | Rabbit | 1:100 | 9661, Cell Signaling |

**Table 4.** Secondary Antibodies.

| Antibody | Company/Cat. No. |
| --- | --- |
| Cy3-Goat α-Rabbit IgG | Jackson ImmunoResearch/ 111-165-003 |
| Cy3-Goat α-Mouse IgG | Jackson ImmunoResearch/ 115-165-003 |
| Alexa Fluor 488-Goat α-Rabbit IgG | Jackson ImmunoResearch/ 115-545-003 |
| Alexa Fluor 488-Goat α-Mouse IgG | Jackson ImmunoResearch/ 115-545-003 |
| Alexa Fluor® 647-conjugated AffiniPure Goat Anti-rabbit | Jackson ImmunoResearch/ 111-605-003 |
| Peroxidase-conjugated AffiniPure Goat Anti-Mouse | Jackson ImmunoResearch/ 115-035-003 |
| Peroxidase-conjugated AffiniPure Goat Anti-Rabbit | Jackson ImmunoResearch/ 111-035-003 |

### Microscopy and quantitative analysis

Brightfield Microscopy: Brightfield microscopy was performed using an Olympus U-PMTVC microscope (Japan) with Plan objectives (Olympus). Sections were captured using a Moticam 2000 digital camera with Motic Images Plus software or a Pannoramic MIDI II scanner (3DHISTECH) and analyzed with CaseViewer software.

Fluorescence Microscopy: Fluorescence microscopy images were captured using a Zeiss Axioplan fluorescence microscope with Plan-Neofluar objectives (Carl Zeiss), a Nikon Ti-E inverted microscope, and a Pannoramic MIDI II scanner (3DHISTECH), with filter sets for DAPI, FITC, and TRITC.

Confocal microscopy: Sections were captured and analyzed using Scion VisiCapture, the NIS-Elements software package (Nikon), and CaseViewer software.

ZEISS Celldiscoverer 7: For the immunohistochemistry experiments involving numerous groups, a ZEISS Celldiscoverer 7 was used for high-content imaging. This system enabled the efficient collection of statistically relevant data by providing high-quality images and handling large volumes of data effectively. The images were subsequently analyzed using QuPath and ImageJ software.

Total fluorescence intensity of striatal dopaminergic projections was quantified using ImageJ software. The striatum was defined as a single region of interest (ROI) based on TH immunoreactivity, and the integrated density was measured. Background fluorescence was determined from an adjacent cortical ROI lacking TH signal. Striatal fluorescence intensity was expressed as integrated density normalized to background fluorescence (Burgess et al., 2010).

Cell quantification was performed on 14 µm OCT-embedded coronal brain sections (6 slide series). Positive and double-positive cells were quantified in systematically sampled sections within defined anatomical regions. Image analysis and cell counting were performed using QuPath and ImageJ software. Cell numbers were analyzed as raw counts per section and used for statistical comparisons between groups.

### Statistical analysis

A two-tailed, unpaired Student’s t-test was used to rigorously assess the significance of differences in numerical data between independent groups. When a significant result was obtained with a one-way ANOVA, Dunnett’s post hoc test was used to further identify and clarify the specific differences among groups. For longitudinal behavioral data with repeated measures across two timepoints, a two-way mixed-effects model (REML) was employed to assess the main effects of age and genotype, and their interaction, followed by Šídák’s post hoc test for multiple comparisons. This approach ensured a comprehensive analysis of the data, allowing for nuanced interpretations of the statistical results. The data are presented as mean ± standard error of the mean (SEM). SEM also represents the error bars in the graphs. All significant changes and interactions were marked by an asterisk between the bars (* for p<0.05, ** for p<0.01, *** for p<0.001, and **** for p<0.0001). The number of experimental replicates (n) was at least 3.

## Declarations

### Author contributions

A.R. and C.B. conceived and planned the study. A.R. and R.S. performed animal experiments. A.R., R.S, M.M., D.K., I.R., J.K., and C.B. performed behavioural and neuropathology data analysis. A.R and R.S. performed statistical analyses for the behavioural and neuropathological experiment and figure generation using GraphPad Prism (version 10, GraphPad Software, USA). Z.Z. and Z.G.O. performed the human genetic studies. L.L. prepared the genotyping data for the C-OPN cohort. A.R., Z.Z., Z.G.O., R.H.F., and C.B drafted the manuscript. All authors reviewed, revised and approved the final manuscript. Z.G.O., R.H.F., and C.B. secured study funding.

## Acknowledgements

This research has been conducted using the UK Biobank Resource under Application Number 45551. The UKB cohort was accessed using Neurohub (https://www.mcgill.ca/hbhl/neurohub). Data for this article were also obtained from the Accelerating Medicines Partnership® (AMP®) Parkinson’s Disease (AMP PD) Knowledge Platform, release version 4.2. For up-to-date information on the study, visit https://www.amp-pd.org. The AMP® PD program is a public-private partnership managed by the Foundation for the National Institutes of Health and funded by the National Institute of Neurological Disorders and Stroke (NINDS) in partnership with the Food and Drug Administration (FDA), National Institute on Aging (NIA), Aligning Science Across Parkinson’s (ASAP) initiative; Celgene Corporation, a subsidiary of Bristol-Myers Squibb Company; GlaxoSmithKline plc (GSK); The Michael J. Fox Foundation for Parkinson’s Research (MJFF); AbbVie Inc.; Pfizer Inc.; Sanofi US Services Inc.; and Verily Life Sciences LLC.

We thank Alexandra Tsitrina from The Research Support Laboratories of Ilse Katz Institute for Nano-Science and Technology at Ben-Gurion University for excellent support in image analysis.

## Data Availability

PD risk GWAS summary statistics are publicly available from the NDPK portal (https://ndkp.hugeamp.org/research.html?pageid=a2f_downloads_280). PD age at onset summary statistic is publicly available on the NHGRI-EBI GWAS Catalog (https://www.ebi.ac.uk/gwas/) under accession number GCST007780.

## Code Availability

The code for all analyses is publicly available on https://github.com/MerZ03/BMP-pathway-analysis.

