## Supplementary Figures for "Bone Morphogenetic Protein Pathway Modulates Parkinson’s Disease Genetic Risk and Promotes Motor Recovery"

**Supplementary Fig. 1**

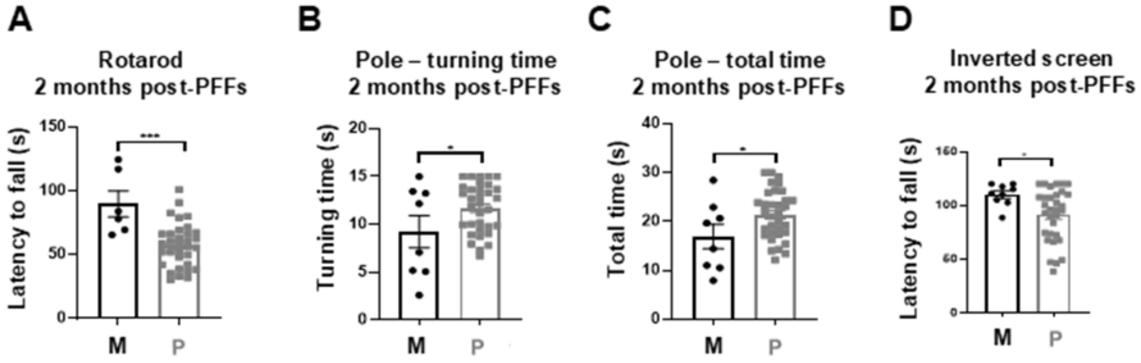


**(A-D)** Rotarod test, pole test and inverted screen test performance is reduced 2 months after α-synuclein PFFs injections. For panels A–D, unpaired two-tailed t-tests were used to compare monomer- versus PFF-injected groups. Significance levels are indicated as: *p < 0.05, **p < 0.01, ***p < 0.001.

**Supplementary Fig. 2**

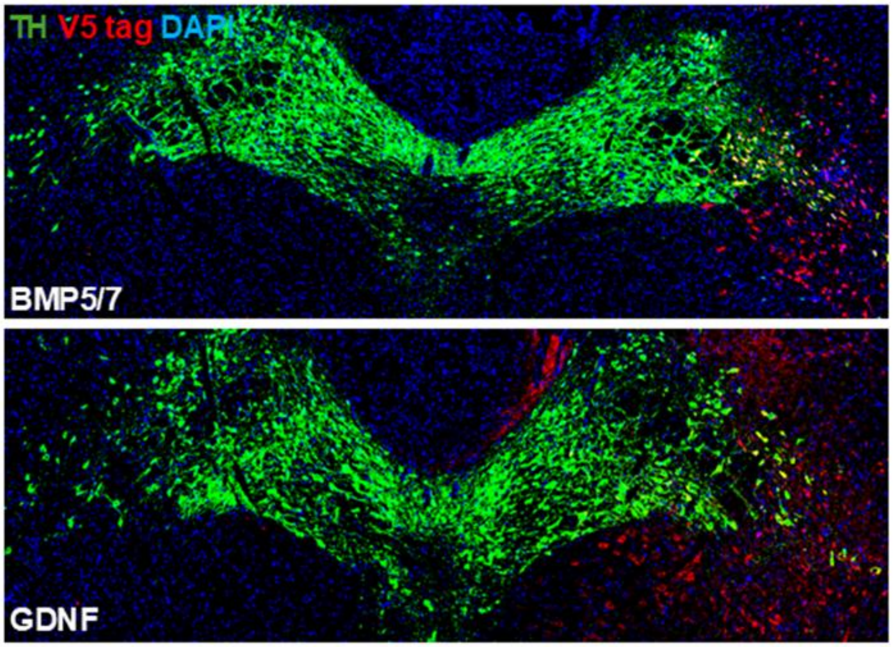


Immunostaining of the SNpc region. V5 tag was used to verify that viral expression of BMP5/7 and GDNF are localized in the SNpc following stereotactic injection.
